# Non-Mendelian inheritance of SNP markers reveals extensive chromosomal translocations in dioecious hops (*Humulus lupulus* L.)

**DOI:** 10.1101/069849

**Authors:** Dong Zhang, Nicholi J. Pitra, Mark C. Coles, Edward S. Buckler, Paul D. Matthews

## Abstract

Genome-wide meiotic recombination structures, sex chromosomes, and candidate genes for sex determination were discovered among *Humulus* spp. by application of a novel, high-density molecular marker system: ~1.2M single nucleotide polymorphisms (SNPs) were profiled with genotyping-by-sequencing (GBS) among 4512 worldwide accessions, including 4396 cultivars and landraces and 116 wild accessions of hops. Pre-qualified GBS markers were validated by inferences on families, population structures and phylogeny. Candidate genes discovered for several traits, including sex and drought stress-resistance, demonstrate the quality and utility of GBS SNPs for genome-wide association studies (GWAS) and Fst analysis in hops. Most importantly, pseudo-testcross mappings in F1 families delineated non-random linkage of Mendelian and non-Mendelian markers: structures that are indicative of unusual meiotic events which may have driven the evolution and cultivation of hops.

## Introduction

The Cannabaceae family of flowering plants has a rich history of contributions to humanity, with the promise of still greater contributions as result new commercial values and invigorated research in two members, *Humulus lupulus* (hop) (2n = 20) and *Cannabis sativa* (hemp, marijuana) (2n = 20) (van Bakel et al., 2011), which diverged around 27.8 Myr (Laursen, 2015). Hop (*H. lupulus*) is a high-climbing, herbaceous perennial, dioecious vine, and has a long history of use as flavoring and stability agent in beer as well as nutraceutical medicine, bio-fuel fermentations and animal fodder (Siragusa et al., 2008). For example, studies of specific hop-derived prenylflavonoids in prevention of cancer, dyslipidemia, and postmenopausal systems spawn interest in metabolic engineering and marker-directed breeding in hop (Ososki and Kennelly, 2003; Stevens and Page, 2004; Nagel et al., 2008; Miranda et al., 2016). The properties of its reproductive system, such as dioecy and obligate outcrossing, high heterozygosity and a large genome size (~2.6Gb), and complex sex-determination system (Neve, 1958), render challenges of genetic dissection of complex traits in hops.

Wild *H. lupulus* is represented by at least five extant species: (1) var. *lupulus* for European wild hops; (2) var. *cordifolius* mainly distributed in Japan, and vars. (3) *neomexicanus* (in the Southwestern U.S.), (4) *pubescens* (in the Eastern/Midwestern U.S.) and (5) *lupuloides* (throughout the northern Great Plains); spreading throughout North America. Asian and North American wild hops resemble each other morphologically, suggesting a genetically closer relationship, while they differ more so from European hops (Murakami et al., 2006). Many contemporary cultivars are hybrids of North American and European genetic materials, in which North American hops have been characterized by their higher bitterness and aroma (Reeves and Richards, 2011) than European cultivars. In other crops, breeding programs have successfully exploited novel genetic variations from wild exotic germplasms into modern cultivars (Tanksley and McCouch, 1997; Bradshaw, 2016) to gain desirable traits such as favored flavors, drought tolerance, and disease resistance. Successes with wild resources and predictions of climate change have spurred resurgence in conservation biology of plant genetic resources (Castañeda-Álvarez et al., 2016; Gruber, 2016).

Molecular marker systems have been developed for hops and applied in genetic mapping of families (reviewed in Henning et al., 2015), including non-referenced GBS markers (Matthews et al., 2013) and GWAS applied to disease resistance (Henning et al., 2015) and sex determination (Hill et al., 2016), using whole genome-referenced GBS markers. However, genetic phenomenon in hops still is under-explained. For example: (1) significant segregation distortion from Mendelian segregation expectations have been repeatedly reported in mapping populations, indicating that the segregation bias was due to genetic properties rather than genotyping errors (Seefelder et al., 2000; McAdam et al., 2013); (2) Unusual female-biased sex ratios have been observed in controlled crosses, where high pollen loads were applied (Jakse et al., 2008). In other genetic systems, segregation distortion is a result of various patterns of meiotic drive and chromosomal (re)arrangements (Taylor and Ingvarsson, 2003). Examples of meiotic drive include the B chromosomes observed in insects (Fontana and Vickery, 1973), the X-linked meiotic drive in Drosophila species (Lyttle, 1993) and the t-haplotype in mice (Silver, 1993). Some well-known examples of chromosomal (re)arrangements in plants include the neocentromeres (knobs) of maize (Buckler et al., 1999), the translocation heterozygosity in some species of the genus *Clarkia* (Snow, 1960), *Oenothera* (Rauwolf et al., 2008; Golczyk et al., 2014) and *Viscum* (mistletoe) (Wiens and Barlow, 1975; Rauwolf et al., 2008).

One classical cytogenetic analysis (Sinotô, 1929) suggested that at least 4 chromosomes in the male of var. *cordifolius* are involved in reciprocal translocation, detected by a chromosome chain in a shape of ‘zigzag’ during the first meiotic metaphase. Our characterization with an unprecedented system of high density genome-wide SNP markers allows detailed examination of translocation heterozygosity, which predisposes rediscovery of the mode of inheritance in this species. Cytogenetic investigations in *Clarkia* (Snow, 1960) and *Oenothera* (Rauwolf et al., 2008; Golczyk et al., 2014) may provide comparative insight on putative translocation heterozygosity in *Humulus*.

However, except for Japanese wild hops (var. *cordifolius*), heterozygotes in males were reported neither in European wild types (var. *lupulus*) nor in North American wild types (var. *neomexicanus*) (Winge, 1932; Jacobsen, 1957; Shephard and Parker, 2000). To date, size ratio of sex chromosomes in wild hops is not fully explained: a description of sex-chromosome variability across subspecies in hops is widely accepted (Shephard and Parker, 2000). Moreover, lack of cytogenetic studies in modern hybrids confuses the mode of genetic inheritance in hops. New insight into genetic parameters in *Humulus* is herein offered by a next generation sequencing (NGS) platform applied to an unprecedented 4512 accessions, including 22 sibling and halfsibling families, genotyped with GBS SNP marker system, comprising 1,235,148 SNPs. Previously reported NGS GBS studies in hop (Matthews, 2013, Henning et al., 2015; Hill et al., 2016) have focused on smaller (511) association panels and reduced marker sets, have not addressed population structure. Furthermore, filtering against SNPs in segregation distortion (SD), thus, ignoring rather than characterizing chromosome regions in SD, resulted in a low-power genetic system that dubiously supported candidate gene discoveries. We present a qualified, high-density marker system applied across a structured association panel and 22 families, analysis of population structure among exotic vs. cultivated accessions within the panel, and a set of strongly supported candidates associated with sex determination. Inclusion, rather than ignorance, of markers in strong SD, has led to testable hypotheses of chromosome structure and recombination constraints in hop, which requires re-assessment of breeding strategies. The need for new, classical cytogenetic studies is implicated by NGS exploration.

## Results

### Phylogenetic relationships of modern cultivars and North American indigenous exotics

European var. *lupulus* is the ancestor of most commercial hops used today, thereby commercial cultivars retain a large proportion of var. *lupulus* genome. In addition, the genetic diversity of hop crop has been contributed by mostly male donors from North America and Asia. To understand the phylogenetic relatedness in hop races, we focused on a subset of 251 accessions, consisting of 183 modern cultivars (CV) including all progenitors of F1 families in this study and 68 wild hops. The neighbor-joining tree (Figure 1a) shows three distinct clusters. The modern cultivars were clustered together, indicating a common derivation in domestication of hops. The other two clusters reflect geographical origins of North American wild hops (Figure 1b), in which one group (SW_wild) includes 22 Southwestern U.S. wild hops (represented by var. *neomexicanus*), and the other group contains 20 wild hops (represented by var. *lupuloides*) from Northern U.S./Canada (N_wild) and 3 (represented by var. *pubescens*) from Midwestern U.S. (MW_wild). Seven wild individuals from Kazakhstan are intermediate among the modern cultivars, consistent with a previous inference (Murakami et al., 2006) of a close genetic relationship between wild hops from Europe and the Altai region (close to western China, located on boundaries of Russia, Mongolia, Kazakhstan and China).

**Figure 1.**
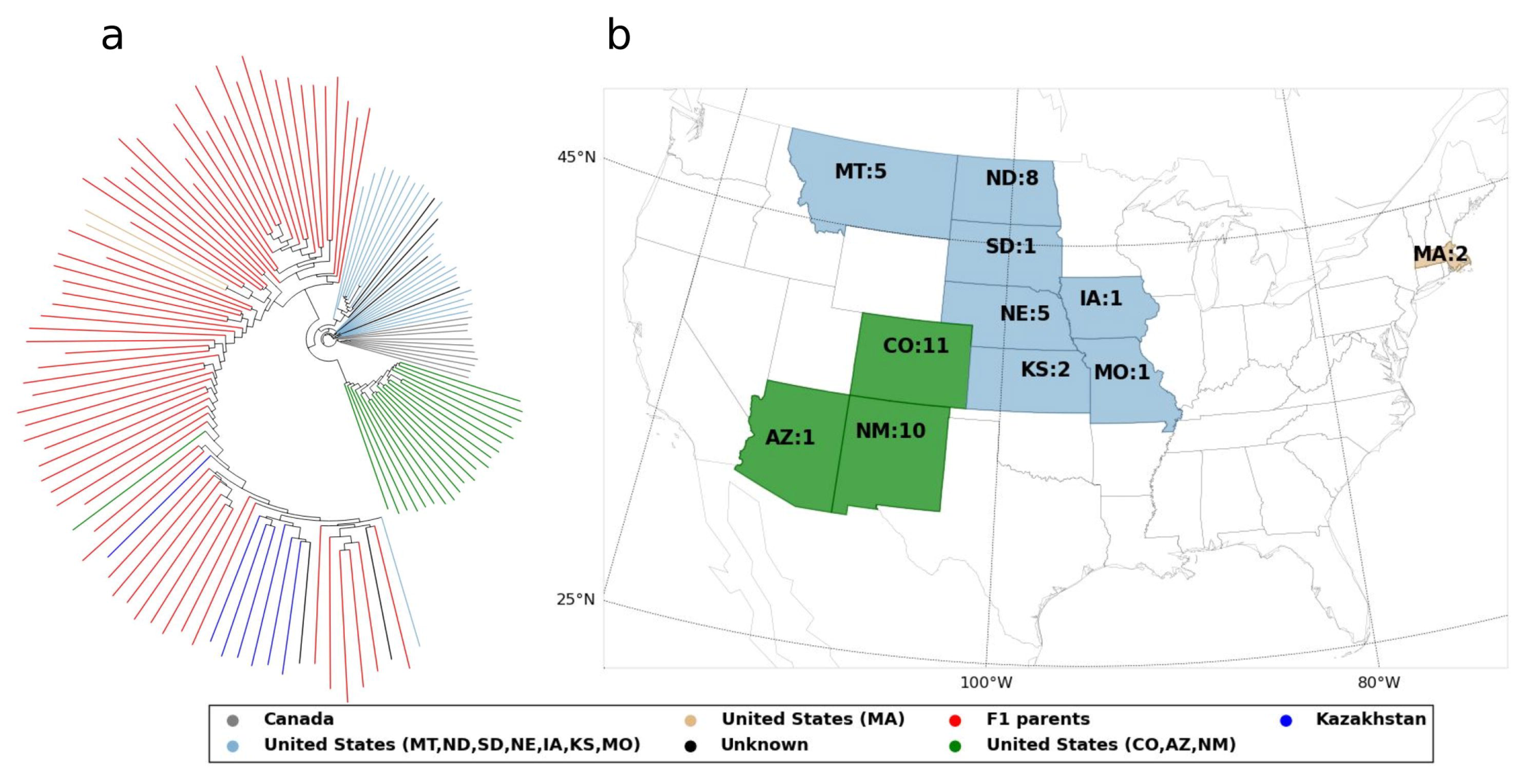
Population structure of 251 hop accessions and geographic origins of the U.S. wild types. 183 modern cultivars are indicated by red color. 68 wild hops are color-coded by geographic origins. (a) Neighbor-joining tree of the 251 hop accessions. (b) The state names are followed by sample counts. Three state groups (“MT, ND, SD, NE, IA, KS, MO”, “CO, AZ, NM” and “MA”) are color-coded to distinguish from one another.

The level of population differentiation, fixation index (Fst), was measured across the three clusters. SW_wild exhibits relatively close genetic relationship (Fst = 0.1663) with N_wild, apparently supporting relatively close ancestry and geographical origins of the two wild populations. Genetic distinction between the modern cultivars and the North American wild hops is evident: [Fst (CV vs. SW_wild) = 0.31; Fst (CV vs. N_wild) = 0.295].

To demonstrate the population structure of F1 families (N ≥ 60) (Figure 2a) in our dataset, we used a nonlinear algorithm (implemented in Python scikit-learn), t-Distributed Stochastic Neighbor Embedding (t-SNE) (Maaten and Hinton, 2008), for dimension reduction of the IBS-based distance matrix. The F1 families derived from genetically divergent progenitors can be easily distinguished from one another, while the half-sibling families exhibit ambiguous clustering patterns. A network of pedigree (Figure 2b) reflects that the F1 families in the current dataset were mostly derived from genetically related varieties.

**Figure 2.**
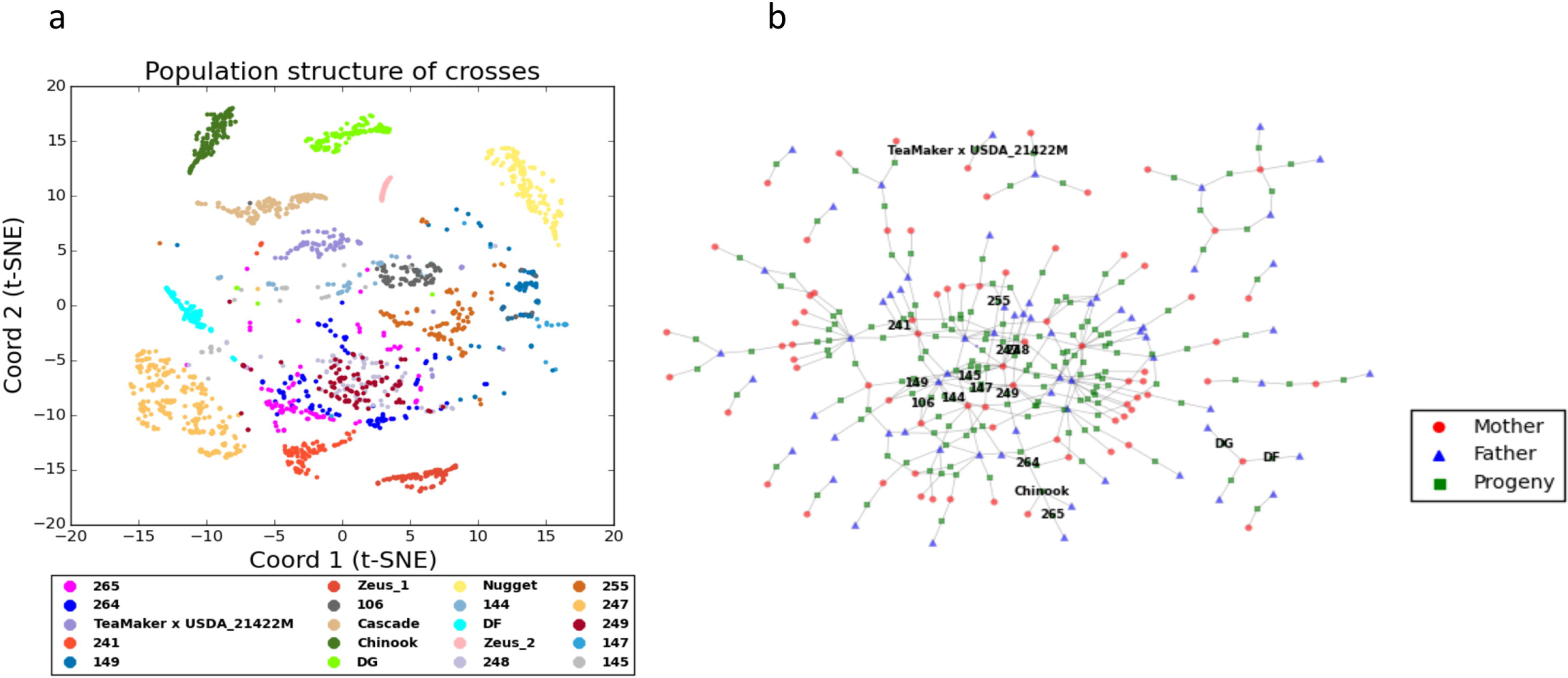
Population structure and pedigree network of F1 families. (a) t-SNE plot for 20 F1 families (N ≥ 60). (b) The overview of pedigree for genotyped F1 families.

### Segregation distortion in hybrids

Ubiquitous presence of non-Mendelian factors results from inherent mechanisms in *H. lupulus* or from genotyping errors. While genotyping errors are random, the genuinely distorted markers can exhibit pronounced correlation with Mendelian segregation markers. On the basis of clustering of pairwise Spearman’s correlation in pseudo-testcross (Pt) markers in three F1 families, we hypothesize that (1) severe SD tends to occur near breakpoints to favor translocation complexes; (2) patterns of linkage can differ across the three populations; (3) a large scale, perhaps genome-wide, meiotic chromosomal complex might occur in the progenitors of the three populations; and (4) translocation heterozygosity may be a ubiquitous phenomenon in hybrids of hops, implicating its genetics and biological significance throughout the cultivation history of hops.

Pseudo-testcross in families “144” and “247” shows multiple ‘super’ linkage groups in terms of their size and inter-marker correlation (Figure 3a,S2a). In family “265”, linkage groups tend to have equal size (Figure S2b), but exhibit relatively high correlation to one another. Alignments across the three sets of clusters (before phasing coupling groups) (Figure 3b,3c) show most of anchor (common) markers were distinctly clustered. Using a nonlinear algorithm (implemented in Python scikit-learn), locally linear embedding (LLE) (Roweis et al., 2000), the consistent clustering patterns (Figure 4) were implicated by projection of genetic maps into spatial coordinates. Moreover, we observed that the Louvain method performs extremely well to phase groups in coupling. The effectiveness of the clustering methods indicates the correlation across linkage groups was caused by real meiotic events.

**Figure 3.**
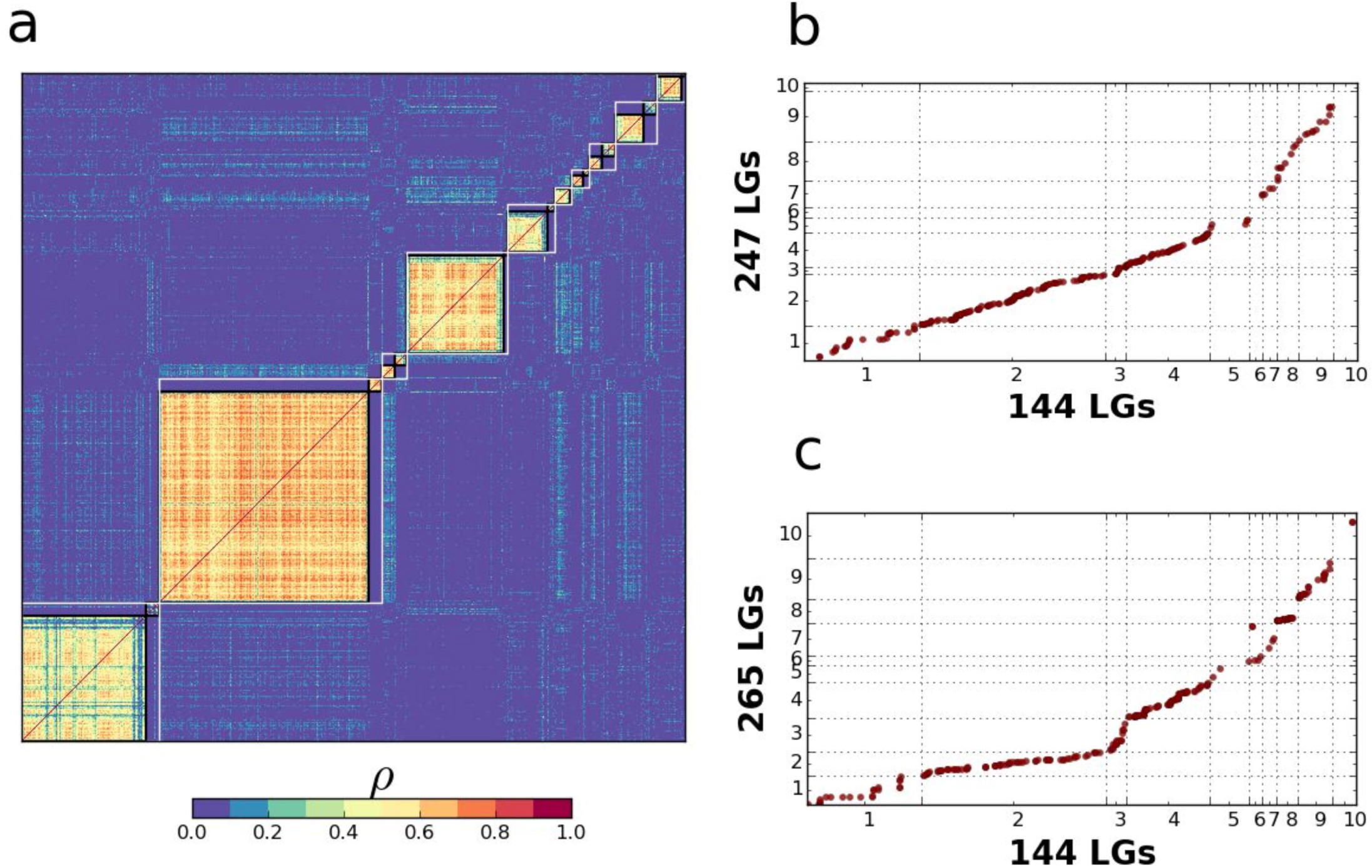
Linkage groups for the maternal line of family “144” and correspondence across 3 genetic map sets. The degrees of Spearman’s correlation (rho) are color-coded. (a) Unphased and phased (linkage for grandparents) groups are bounded by white and black frames individually. (b) Alignment of unphased groups between “144” and “247”. (c) Alignment of unphased groups between “144” and “265”.

The loci with 5% ≤ MAF < 15%, deviated significantly from the 1:3 expectation for Pt markers, account for 28.3%, 49% and 48.3% in families “144”, “247” and “265” respectively, in which proportions of the distorted loci correlated (rho ≥ 0.3) to the Mendelian segregation markers (15% ≤ MAF ≤ 35%) are 78.3%, 48.9% and 71.8%.

Spatial coordinates of Pt markers, in accordance with correlation heatmaps, offer a complete picture (Figure 4) of genetic linkage with and without inclusion of SD markers, which appear to play a role in bridging Mendelian markers. Such linkage patterns were constantly observed in the three families (Figure 4, S3).

**Figure 4.**
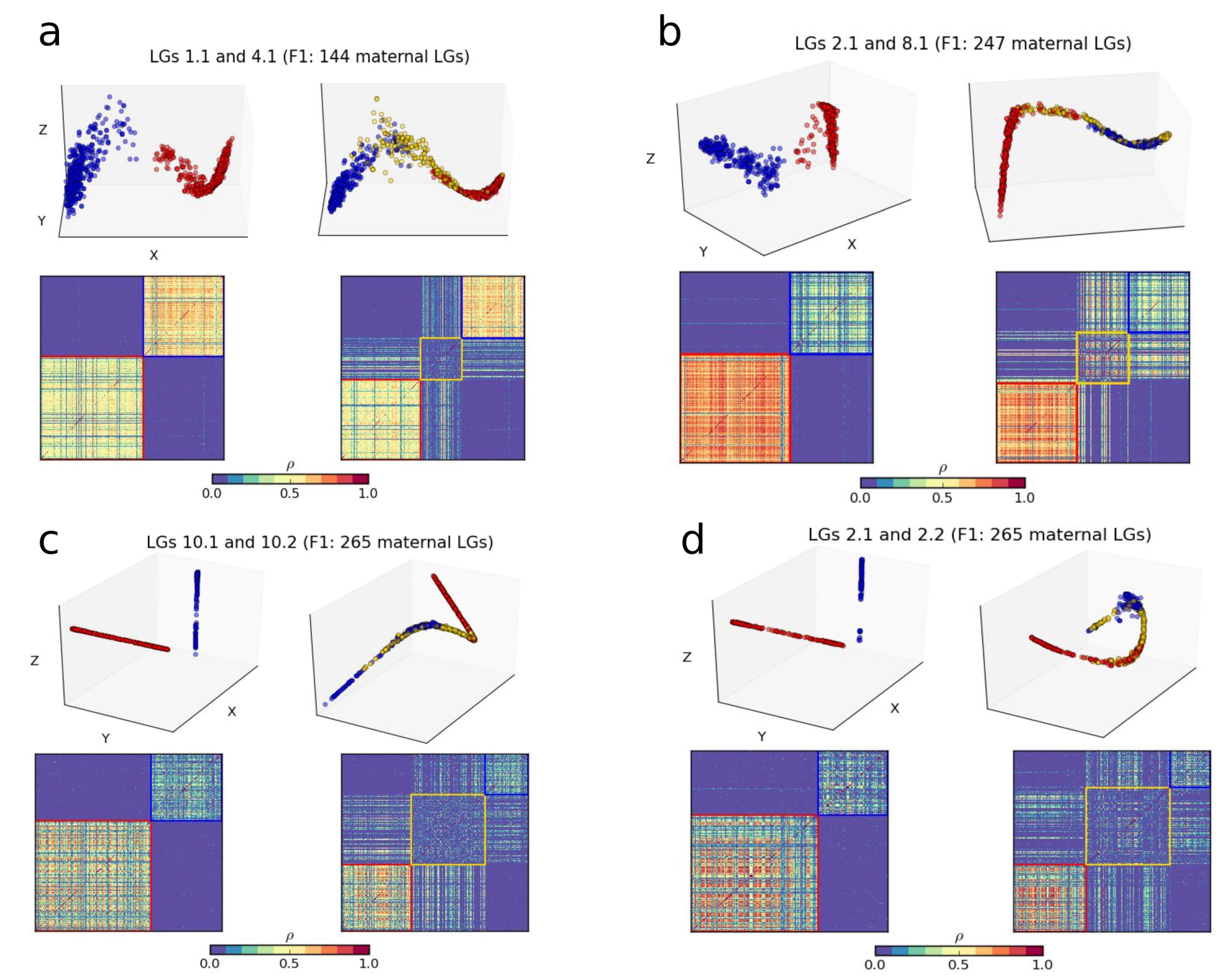
Linkage of Mendelian (15% ≤ MAF ≤ 30%) and non-Mendelian Pt markers (5% ≤ MAF < 15%), based on Spearman’s correlation (rho). In each sub-figure, clustering patterns without (left) and with (right) inclusion of segregation distortion are presented by LLE (top) and the Louvain Modularity (bottom). Mendelian markers in two linkage groups are indicated by blue and red colors individually. Segregation distortion (SD) markers are indicated by yellow color.

We show the correlation across the 5 largest linkage groups (Figure 5a) in family “265” based on the spatial representation of the positions of the markers with 15% ≤ MAF ≤ 35%. The model (Figure 5b) clearly depicts distortion decreases, when the markers are more distal to the convergence areas, and indicates that the progenitors of family “265” experienced translocation to form a large chromosomal complex.

**Figure 5.**
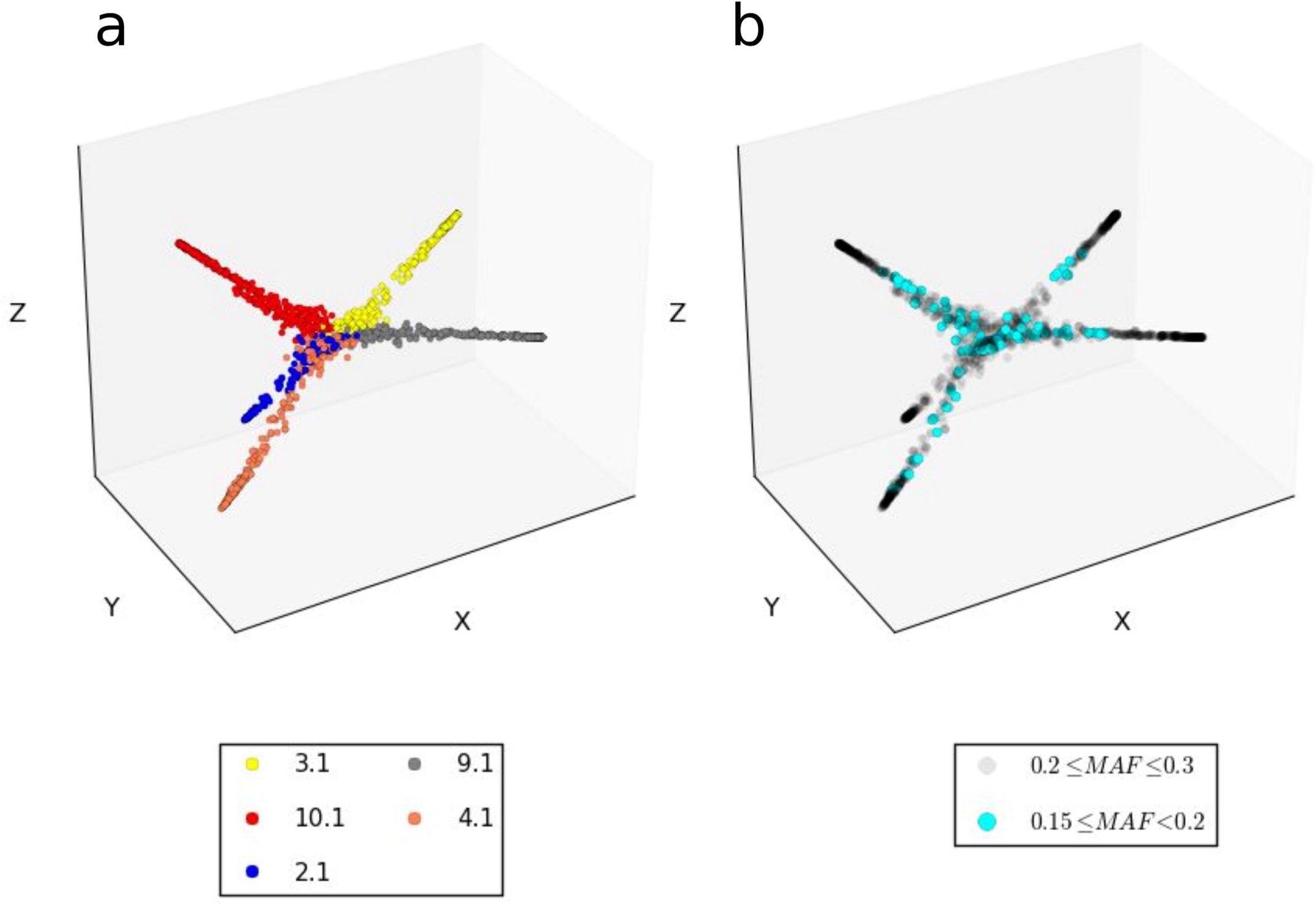
Linkage patterns of the 5 largest linkage groups in family “265”, based on spatial coordinates defined by LLE. (a) Linkage groups are color-coded. (b) Markers with 0.15 ≤ MAF < 0.2 and 0.2 ≤ MAF ≤ 0.3 are distinguished by cyan and gray colors.

One linkage group (LG) in one family corresponding to multiple groups in the other family, would suggest corresponding loci were involved in chromosomal rearrangement in progenitor of the former family. One striking case (Figure 6) can be found in LG2.1 in “144” corresponding to two coupling LGs (2.1 and 2.2) in “265”. Two control correspondences (LG1.1-LG1.2 and LG3.1-LG3.1) were used to illustrate the power of the clustering approaches. However, such one-to-multiple correspondence was seldom observed across the three families. That may reflect the conservation of chromosomes positioning in the heterozygotes complex and invariable occurrence of the translocation heterozygotes in the progenitors of the three families.

**Figure 6.**
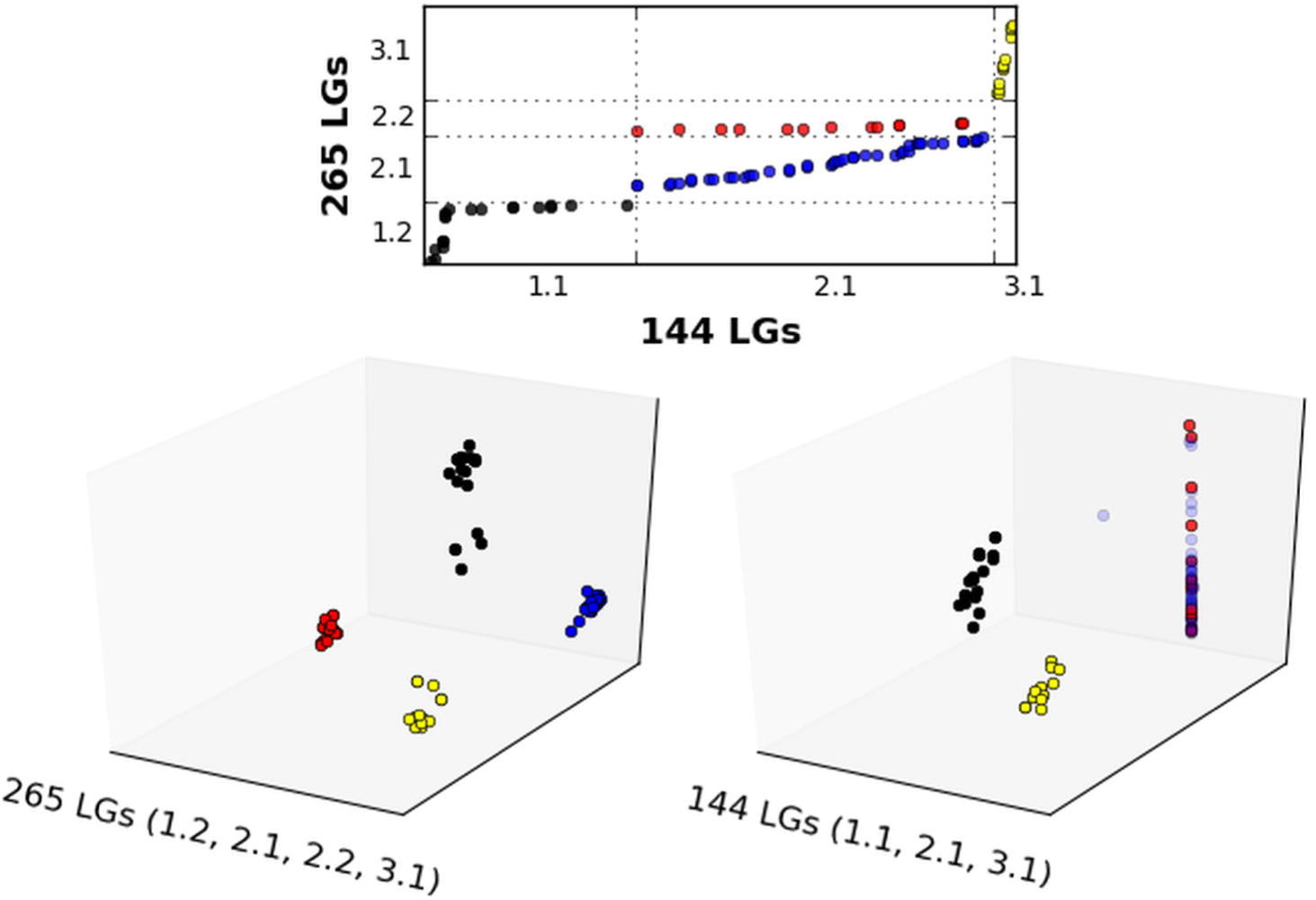
One-to-two genetic correspondence between “144” and “265”. LG2.1 in “144” corresponds to LG2.1 and LG2.2 in “265” (top sub-figure). Two instances of one-to-one correspondence (LG1.1-LG1.2 and LG3.1-LG3.1) are added for control. Spatial representations of linkage groups (bottom sub-figure) in the two families were derived from LLE.

### GWAS for sex determination

Sexual phenotype regulation is a particularly important problem in dioecious plants, herein exemplified by hop (*H. lupulus*). Male and female hop flowers can be easily distinguished; the male flower closely resembles a typical perfect flower, while female inflorescence meristems produce flowers arranged in ‘cones’ (Shephard and Parker, 2000).

We used a mixed linear model to assess evidence of phenotype-genotype association. In families “247” (N = 364, N_male_ = 30) and “265” (N = 95, N_male_ = 13), linkage group (LG) 4 (Figure 7a,S4) consistently shows the most striking association with sex, even though “265” has a small effective population size. This signal was additionally supported by Fst mapping in “247” (Figure 7b). However, pseudo-testcross accounts for part of association signals. To perform genome-wide scan, we assessed association between 356,527 markers and 850 individuals (N_male_ = 129, N_female_ = 721), as described in Methods.

**Figure 7.**
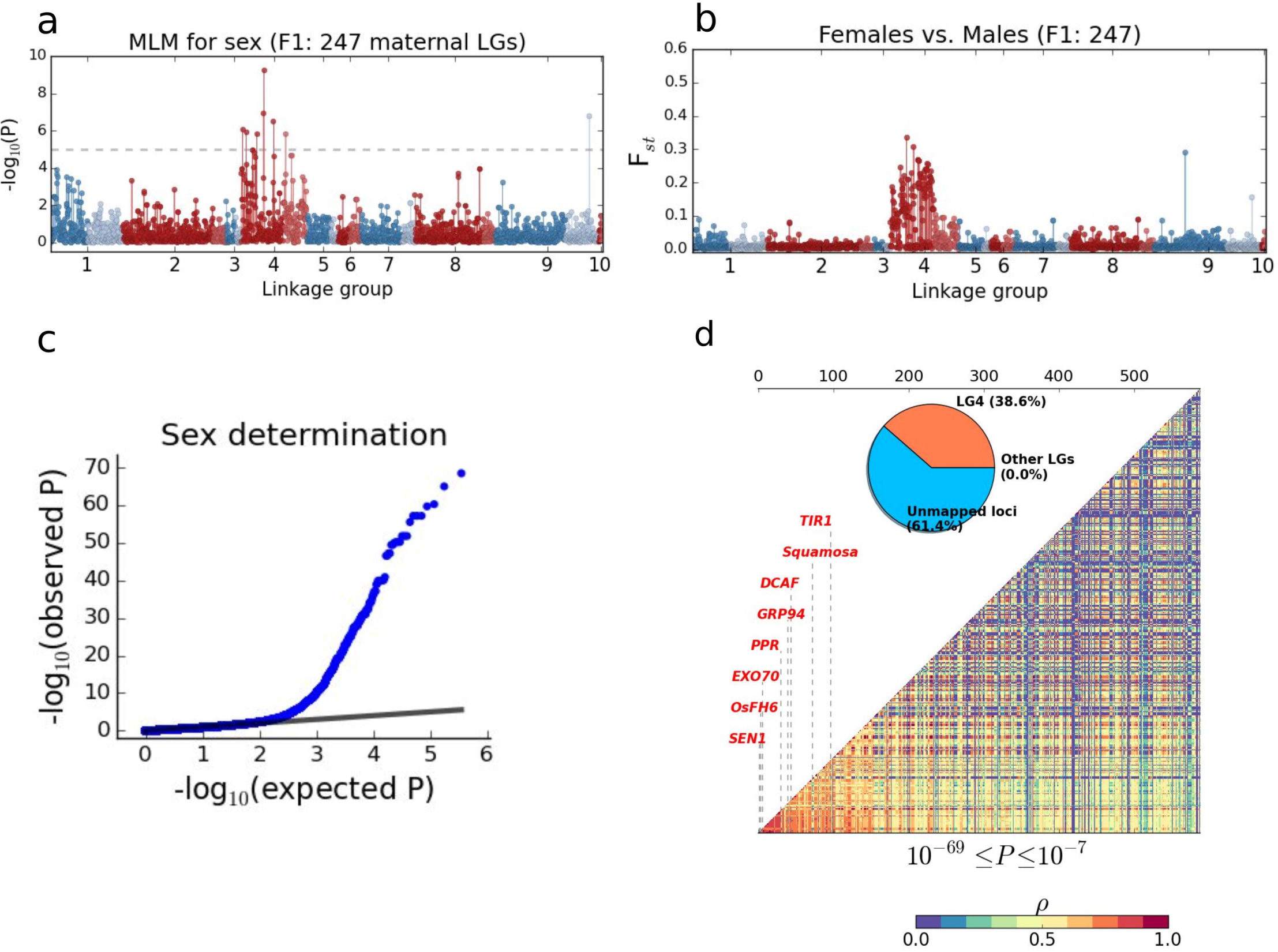
Association studies and Fst mapping of sex determination in hop. (a) Linkage group-based Manhattan-plot of MLM for sex determination in family “247” (N = 364, N_male_ = 30). Light and deep colors are used to distinguish two phases (linkage for grandparents) in coupling. (b) Manhattan-plot of Fst in females vs. males in “247”. (c) Log Quantile-Quantile (QQ) plot of 356,526 association tests (SNPs) for sex determination in 850 individuals (N_male_ = 129, N_female_ = 721). (d) Correlation among 588 association (P ≤ 10^−7^) markers, the proportions of 588 markers in LG4, other LGs and unmapped data set, and 8 gene candidates for sex determination in hop.

A total of 588 SNPs with P ≤ 10^−7^ were identified (Figure 7c,7d). We noted that LG4 and other LGs respectively account for 38.6% and 0.0% of the association markers, reinforcing the importance of LG4 in sex expression in hops. On the basis of pairwise Spearman’s rho, we observed high correlation among the 588 SNPs, implicating that the association markers mostly come from one linkage disequilibrium (LD) block. Adding up scaffolds showing association approximates ~9.75Mb of the mapping resolution accounting for ~0.38% of the hop genome.

To identify gene candidates for hop’s sex, we aligned by BLAST the top 100 scaffolds against UniProt and NCBI protein databases to search for genes with known function involving in the mechanism of sex determination (Table S4). A total of 8 gene candidates (Ruegger et al., 1998; Ishiguro et al., 2002; Koizuka et al., 2003; Hála et al., 2008; Shimizu et al., 2008; Zhang et al., 2008; Chen et al., 2010; Zhang et al., 2011) were obtained (Figure 7d), in which 7 blast hits have 81%-99% length matching to the target proteins and 7 hits encompass ≥ 1 the significant association site (p ≤ 10^−25^).

In hop, a significant deformation of the apical meristem, the producer of flower primordia cells, has been observed during the transition to the reproductive phase, resulting in the apparent morphological differences between male and female flowers (Shephard and Parker, 2000). Two notable examples, relating to the floral structures, are (1) a glucose-regulated protein 94 (GRP94)-like protein on scaffold LD152823 that is known in Arabidopsis affecting shoot apical meristems, floral meristems and pollen tube elongation (Ishiguro et al., 2002); and (2) a Squamosa-like protein, identified on scaffold LD147778, has essential roles in vegetative phase change and flower development in multiple plants (Chen et al., 2010).

### Genetic differences and phenotypic variation across populations

To assess genetic contributions to between-population phenotypic differences, we used Fst analysis to characterize genetic variations across var. *neomexicanus*, var. *lupuloides* and CV. Previously cloned genes in *Humulus* were highlighted to suggest that with respect to essential chemical composition and drought tolerance, hotspots with unusually high or low Fst values deserve a great deal of attention.

A linkage map-based view of Fst highlights two notable patterns (Figure 8). First, the degree of genetic variation, as expected, is much greater in CV vs. *neomexicanus/lupuloides* than in *neomexicanus* vs. *lupuloides*. Regions that exhibit above-average population differentiation in CV vs. *neomexicanus* typically also exhibit above-average population differentiation in CV vs. *lupuloides*. Second, the 5 largest linkage groups account for a large proportion of genetic variation between populations. The 90^th^ percentile of Fst values (Figure 8d), referred to as 0.63 in CV vs. *neomexicanus* and 0.62 in CV vs. *lupuloides*, were used to define the significant genetic difference.

**Figure 8.**
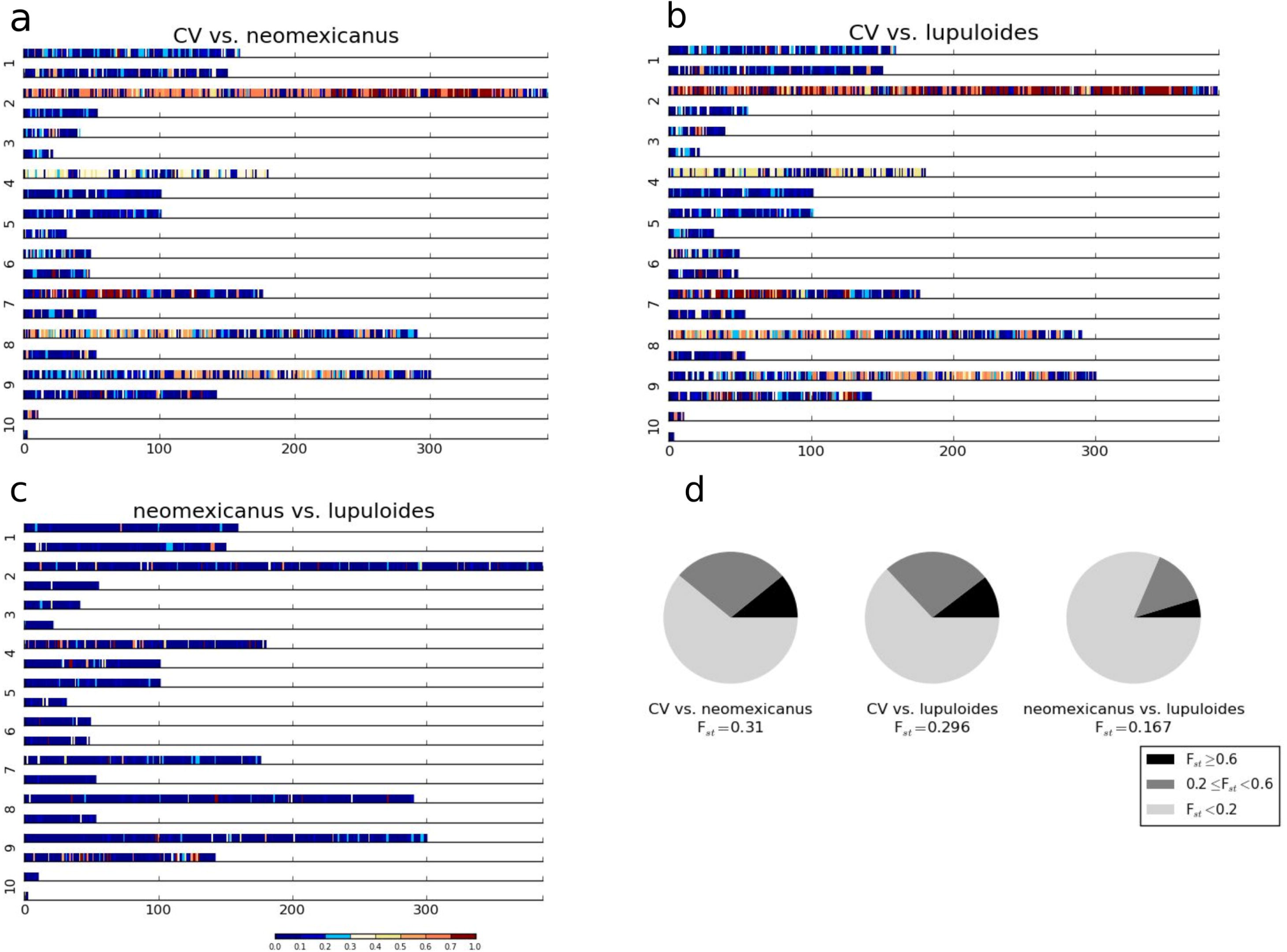
Linkage group (in family “247”)-based Fst heatmaps and the overall Fst distribution. Population differentiation (a) between modern cultivars (CV) and var. *neomexicanus*; (b) between CV and var. *lupuloides*; (c) between var. *neomexicanus* and var. *lupuloides*. (d) Spectrum of the overall Fst distribution.

In beer brewing, the chemical composition, resins imparting bitterness and essential oils contributing to flavor and aroma, determine the quality and flavor of hops. The key brewing resins include alpha-acids and beta-acids, also referred to as humulones and lupulones respectively. It is generally accepted that North American hops have high content of alpha acids, rendering bitter beers, while European hops have lower content of resins and essential oil. Enhancement of alpha acids content in the contemporary cultivars was accomplished by crossing European hops (var. *lupulus*) to North American donors, one of which resembles var. *lupuloides* from Canada (Neve, 1991; Reeves and Richards, 2011). Genes affecting resins and essential oil can be found in the regions presenting genetic divergence between North American hops and adapted varieties.

In hop, two homologues of chalcone synthase (*CHS*) in bitter acid biosynthesis have been cloned (Figure S5), referred to as *CHS_VPS* (GenBank: AB015430.1) (Okada and Ito, 2001) and *CHS_H1* (GenBank: CAC19808.1) (Matoušek et al., 2002) respectively. Both genes are highly active in lupulin glands, to serve as catalysts of the synthesis reactions of alpha-acid and beta-acid. Significant genetic differences (Fst ≥ 0.6) between cultivars and North American wild types emerge in a region adjacent to (~40Kb away from) *CHS_H1*. In contrast, a surrounding region (~15Kb away from) of *CHS_VPS* harbors moderate Fst (= ~0.4) in CV vs. *neomexicanus*, and low Fst (= ~0.1) in CV vs. *lupuloides*. The differences of population differentiation in the surrounding regions of the two *CHS* homologues may reflect genetic introgression and differential allele selection resulting from domestication towards higher alpha acid yields.

Var. *neomexicanus* is traditionally grown in rain-fed or supplementary irrigated areas, and has capability to withstand periods of water deficit and to yield an economic return to farmers. In contrast, hops frequently occurring in the high latitude of the northern North America favor plenty of water and sunlight. Likewise, heavy irrigation is required in modern hop crops. Hence, genic regions exhibiting the significant population differentiation in CV vs. *neomexicanus*, but not in CV vs. *lupuloides* can be prioritized to identify gene candidates affecting drought tolerance.

Two intriguing proteins were identified on scaffold “LD161390” (Figure 9a), which are a *H. lupulus* basic leucine zipper transcription factor Long Hypocotyl 5 (*bZIP HY5*) (GenBank: CBY88800.1) (Matousek et al., 2010) and a light-harvesting chlorophyll (*LHCII*) a/b binding protein (*LHCB*) (NCBI RefSeq: XP_002307004.1).

**Figure 9.**
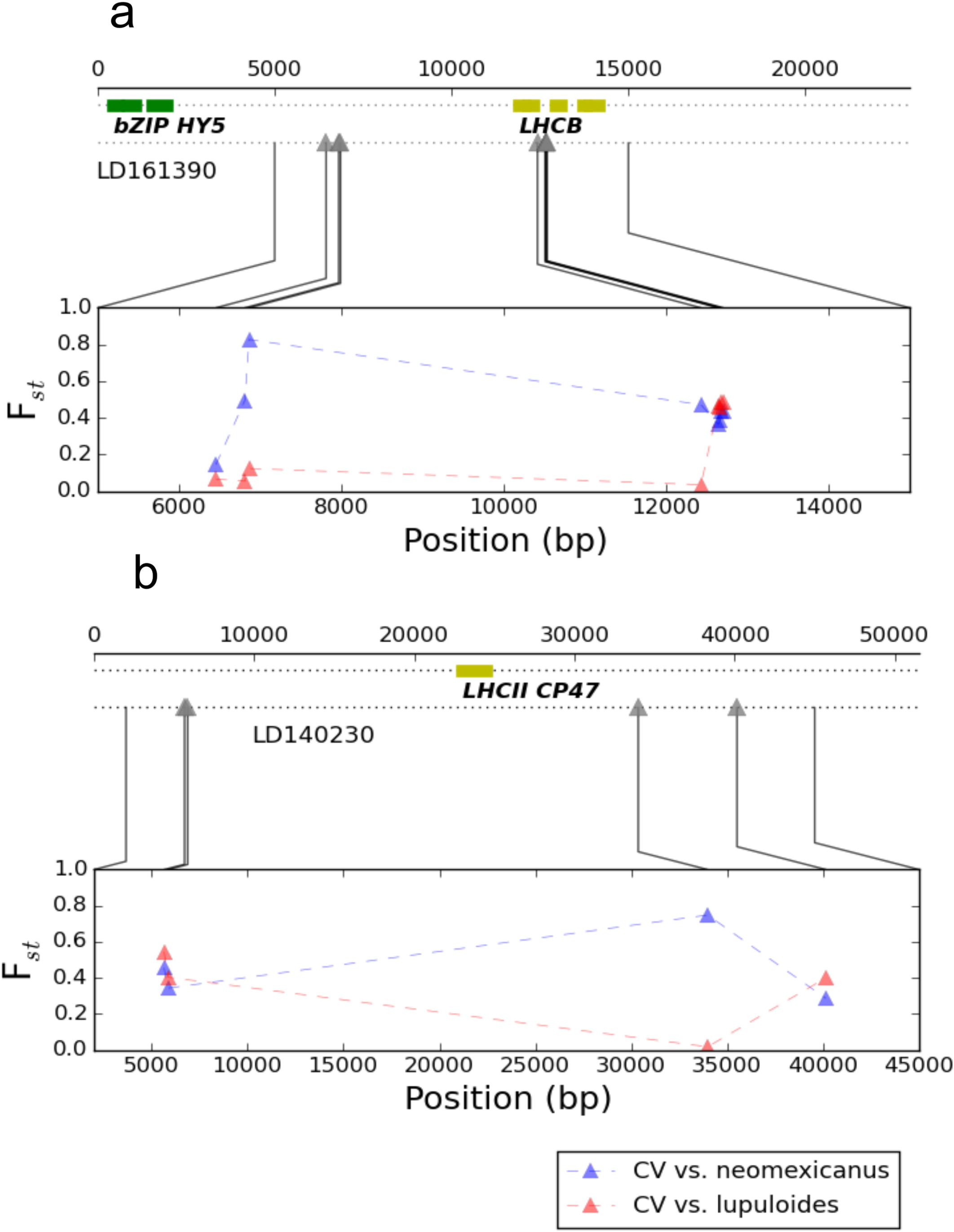
Three gene candidates for drought tolerance in hops. Between-population Fst values are indicated on the scaffolds where the three candidates are located. (a) The approximate positions of *bZIP HY5* (GenBank: CBY88800.1) (Matousek et al., 2010) and *LHCB* (NCBI RefSeq: XP_002307004.1) on scaffold “LD161390”. (b) The approximate position of *CP47* (NCBI RefSeq: YP_009143579.1) on scaffold “LD140230”

*HY5* has a notable relationship with the phytohormone abscisic acid (ABA), a plant hormone in response to environmental stress, such as high salinity, drought and low temperature (Nakagawa H, Ohmiya K, 1996). Via binding and promoting *ABI5*, encoding a bZIP transcription factor, *HY5* can mediate *ABA* sensitivity (Chen et al., 2008) to induce stomatal closure, thus reducing transpiration and preventing water loss from leaves.

The putative *LHCB*, physically close to the *HY5*-like gene, embeds a mutation with striking Fst in CV vs. *neomexicanus*, but not in CV vs. *lupuloides*. *LHCB* involves in the steps of photosynthesis by capturing sunlight and balancing excitation energy between photosystems I (PSI) and II (PSII). It is clear that over-expression of a *LHCB* is a key to enhance stomatal sensitivity to ABA in green alga (*Dunaliella salina*) (Liu and Shen, 2004).

A previously *C. sativa* (marijuana) *LHCII* protein, *CP47* (NCBI RefSeq: YP_009143579.1), is present on scaffold “LD140230” (Figure 9b), near markers showing the pronounced population divergence between *neomexicanus* and CV/*lupuloides*. In sorghum, *CP47* serves as a connection between the main light harvesting complex LHCII and reaction center of PSII. The characteristic adaptation of sorghum to drought may be partly related to downregulation of *CP47* during drought stress and high irradiance (Masojídek et al., 1991).

## Discussion

Hop crop acreage and usage is rapidly expanding and diversifying because of a bourgeoning craft brewing industry. Hop breeding programs have been long attempting to exploit genetic resources for bitter flavor, aroma and disease resistance. However, a worsening drought and unseasonably hot weather may decentralize the targets of breeding programs soon. For example, in Europe and the US, most hop farms experienced severe water shortage in 2015. Like many other crops, exploitation of novel genetic variation in response to drought stress is of paramount importance for a sustainable hop production system.

Understanding genetic recombination is essential for speed and accuracy of plant breeding. Indeed, it is generally difficult to breed new commercial hop varieties through mass selection and crossing. Our findings show that a large scale, perhaps genome-wide, chromosomal rearrangement may occur in the progenitors of F1 families. Translocation heterozygosity can extend linkage to nonhomologous chromosomes, and favor severe segregation distortion accumulated near the translocation breakpoints (Taylor and Ingvarsson, 2003; Farré et al., 2011). Such high degree recombination suppression may hinder effectively selection of desired, novel allele combinations. Inasmuch, breeding strategies favoring introgression of diversity deserve re-emphasis in addition to extended attention to recombination and segregation phenomena in traits. Manipulation of genetic recombination in hops also deserves further focus.

At least 57 species of flowering plants were characterized by permanent translocation heterozygotes (Holsinger and Ellstrand, 1984), in which *Clarkia* (2n = 18) may provide a comparative system to hypothesize meiotic configurations in *Humulus*. Translocation polymorphism has been observed in at least 14 of the 34 known species in *Clarkia* (Snow, 1960). Judging from cytogenetic analyses in 9 natural populations of *Clarkia dudleyana*, none or few translocation heterozygotes occurred within individual colonies, while extensive translocation heterozygotes invariably arose from hybrids derived from cytologically distinct races (Snow, 1960). The largest heterozygotes complex consists of 18 chromosomes. For *H. lupulus* wild types, until now, researchers have still been at odds about the size ratios of sex chromosomes. Except for a tetravalent in var. *cordifolius*, no meiotic chromosome associations were reported in other wild types, suggesting that the five wild *H. lupulus* species are cytologically differentiated. Segregation distortion presenting in cultivated mapping populations may stem from historical hybridization across isolated populations of hops.

Based on the projection of SNPs into a 3D coordinate system using LLE, the nondisjunction of the 5 largest LGs in “265” provides an example of funnel-shaped representation of a heterozygotes complex. Specifically, chromosome segments distal to and close to translocation breakpoints can be represented by wider and narrower end of the funnel-shaped structure respectively. Apparently, two ends of the funnel also characterize clusters of SNPs with similar allele frequencies. There is a need for cytogenetic studies for drawing conclusions regarding the exact meiotic configuration in *Humulus* hybrids for both males and females. Whether the translocation heterozygosity is sex-linked deserves further investigation.

Translocation heterozygosity may have an important connection to the significantly distorted sex-ratio in favor of females in hops. Likewise, female-biased sex ratios have been found in Mistletoe, another notable dioecious case of translocation heterozygosity. Known that, to maintain heterozygosity, *Oenothera*, a notable monoecious case of translocation heterozygosity, utilizes a system of balanced lethal to purge the lethal homozygotes (Steiner, 1956; Harte, 1994), which is referred to as “recessive lethals”. In the context of XY system, heteromorphism of sex chromosomes dictates that males are more severely affected than females by “X-linked recessive lethals”, because males only have one copy of the X chromosome. Hence, *H. lupulus* may use a system of balanced lethals at the expense of male offspring to preserve genetic heterozygosity in hybrids.

Our results are compelling for translocation heterozygosity studies in light of high-density molecular markers in many other biota. For example, such large scale recombination suppression is also presented in at least 10 species of termite, some types of centipede, and perhaps all of the monotremes (Holsinger and Ellstrand, 1984; Rowell, 1987; Rens et al., 2004). Beyond homologous crossover, translocation heterozygosity has shown considerable evolutionary interest and selective advantage in its own right.

## Materials and Methods

### Plant materials

Hops used in this study were grown under standard agronomic conditions at the Golden Gate Ranches, S.S. Steiner, Inc, Yakima, WA. The un-domesticated, exotic hops are from the National Clonal Germplasm Repository in Corvallis, Oregon (accession details in Table S1-S3). Fifty milligrams of young leaf tissues were extracted in a 96 well block using Qiagen Plant DNeasy Kits and was tested for quality, quantity, and purity, prior to library preparations, using a Agilent 2100 Bioanalyzer (Applied Biosystems, Foster City, CA) and Life Technologies (Carlsbad, CA) Qubit 3.0 Fluorometer. The GBS libraries were prepared using the *Ape*K1 enzyme according to Elshire, et al. (Elshire et al., 2011). Pools of 96 accessions were sequenced on one lane of an Illumina HighSeq 2000 (Illumina, San Diego, CA)

### SNP calling and quality control

The reference sequence refers to a draft haploid genome sequence of Shinshu Wase (SW) (Natsume et al., 2015), which is a modern cultivar bred from a seeding selection cross between Saazer and White Vine-OP. The draft genome, with a total size of 2.05 Gb, consists of ~130,000 scaffolds covering approximately 80% of the estimated genome size of hop (2.57 Gb).

Tassel 5 GBS v2 Pipeline (Glaubitz et al., 2014) was applied to identify tags with at least 10x total coverage, and to call SNPs. Tag sequences were mapped to the reference genome using BWA aligner.

One main source of erroneous SNP calling is misalignment caused by incomplete reference genome, gene duplication and low-complexity regions. To filter out erroneous SNPs due to misalignment, we used two criteria: (1) SNPs with an excessive coverage (e.g. read depth > 127) can be false positives. For GBS data, the maximum read depth for one genotype is unlikely to exceed 127 (Glaubitz et al., 2014). Indeed, we observed that heterozygosity rates and MAF are significantly increased when read coverage exceeds 127 (Figure S1). (2) The orientation of paired reads of the cultivar Apollo (unpublished data), a highly used maternal line in our F1 families, was used to detect false positive SNPs caused by gene duplications. Paired-end alignment was generated by BWA Sampe. Identification of correctly aligned regions was based on SAM flags indicating reads mapped in proper pairs. Using criteria (2) was able to detect ~73% SNPs with the excessive coverage.

### Pseudo-testcross

Three F1 families were used to conduct pseudo-testcross (Pt) recombination mappings, including (1) “144” (N = 179) derived from a cross between Nugget (maternal line) and Male50 (paternal line); (2) “247” (N = 364) derived from two parental lines, Super Galena and Male15; (3) “265” (N = 95) derived from a cross between Chinook and Male57. Using markers heterozygous in the maternal line and null in the paternal line, three genetic map sets were constructed, consisting of 3551 SNPs for “144”, 2369 SNPs for “247” and 4506 SNPs for “265”.

Our analyses followed the main steps in HetMappS pipelines (Hyma et al., 2015). Specifically, (1) to remove contaminants, identity by state (IBS) based distance matrices calculated by TASSEL (Bradbury et al., 2007) were used to identify outliers for each family; (2) SNPs having both parental genotypes (e.g. AA×Aa) with read depth ≥ 4 were retained for the next step; (3) in progeny, SNPs with average read depth ≥ 4 and with site coverage ≥ 50% were retained for the next step; (4) to eliminate the effect of under-calling heterozygotes and sequencing errors, we masked progeny genotypes with depth=1, and converted genotypes aa to Aa because genotype aa cannot exist for parental genotypes AA×Aa in Pt; (5) after correction, SNPs with 15% ≤ MAF ≤ 35% were selected to create linkage groups, and SNPs with 5% ≤ MAF < 15% were deemed the pronounced SD markers; (6) to cluster and order markers, an adjacency matrix with Spearman’s correlation (rho) were derived from the remaining SNPs; (7) on the basis of absolute values of rho, the Louvain method (Blondel et al., 2008) implemented in NetworkX (http://networkx.github.io/) was applied to detect communities (clusters). The Louvain method is an efficient algorithm for community detection in large networks. A similar method, modulated modularity clustering (MMC) (Stone and Ayroles, 2009), has been successfully applied to construct linkage groups; (8) to identify coupling phase from each “absolute rho” cluster, negative values of rho were set to zero, and the Louvain method was applied to positive values of rho (Hyma et al., 2015); (9) MSTmap (Wu et al., 2008) was used to provide a sub-optimal solution of genetic ordering within each linkage group.

Putative 10×2 linkage groups in coupling were obtained in each F1 family. Linkage groups may not represent one chromosome due to pseudo-linkage resulting from chromosomal rearrangement, as discussed in Results.

### Genome-wide association studies (GWAS)

An association population includes 850 individuals, in which 837 (116 males and 721 females) are progeny in 6 F1 families and 13 are paternal lines. Male and female were encoded as ‘1’ and ‘0’ individually. A total of 356,527 SNPs with coverage ≥ 50% and MAF ≥ 5% were retained. The Mixed Linear Model (MLM) (Bradbury et al., 2007; Lipka et al., 2012) was used to assess genotype-phenotype association. The Bonferroni method was used to adjust the significance cutoff for an overall probability of 0.05 for type I error.

## Additional files

**Supplementary Figures.** The file contains supplementary figure S1-S5.

**Supplementary Tables.** The file contains supplementary tables. (**Table S1** Pedigrees of genotyped F1 populations. **Table S2** Cultivar and landrace accessions. **Table S3** Wild exotic accessions. **Table S4** BLASTX hits for scaffolds encompassing sex association (P ≤ 10^−10^) SNPs. Eight strongly supported gene candidates are highlighted.).

## Acknowledgments

We thank Buckler lab and Qi Sun’s group at Cornell for helpful discussions. We thank the growers at Golden Gate ranches for cultivation of experimental plants.

